# Clonal evaluation of early onset prostate cancer by expression profiling of ERG, SPINK1, *ETV1*, and *ETV4* on whole mount radical prostatectomy tissue

**DOI:** 10.1101/667832

**Authors:** Zhichun Lu, Sean R. Williamson, Shannon Carskadon, Pavithra D. Arachchige, Gaury Dhamdhere, Daniel S. Schultz, Hans Stricker, James O. Peabody, Wooju Jeong, Dhananjay Chitale, Tarek Bismar, Craig G. Rogers, Mani Menon, Nilesh S. Gupta, Nallasivam Palanisamy

## Abstract

Expression profiles of ETS related genes and SPINK1 in early onset prostate cancer have not been thoroughly explored. We retrieved 151 radical prostatectomy specimens from young men with prostate cancer (<55yrs) and characterized the expression of ERG, SPINK1, *ETV1* and *ETV4* by dual immunohistochemistry and dual RNA *in-situ* hybridization. Age, race, family history, preoperative prostate-specific antigen, biochemical recurrence and pathological variables using whole mount radical prostatectomy tissue were collected. 313 tumor nodules from 151 men including 68 (45%) Caucasians and 61 (40%) African Americans. Positive family history of prostate cancer was observed in 65 (43%) patients. Preoperative prostate-specific antigen ranged from 0.3 to 52.7 ng/ml (mean 7.04). Follow-up period ranged from 1 to 123.7 months (Mean 30.3). Biochemical recurrence was encountered in 8/151 (5%). ERG overexpression was observed in 85/151 (56%) cases, followed by SPINK1 in 61/151 (40%), *ETV1* in 9/149 (6%), and *ETV4* in 4/141 (3%). There were 25/151 (17%) cases showing both ERG and SPINK1 overexpression within different regions of either the same tumor focus or different foci. Higher frequency of ERG overexpression was seen in younger patients (≤ 45 years old) (76% vs. 49%, *p* = 0.002213), Caucasian men (71% vs. 41% *p* = 0.000707), organ-confined tumors (64% vs. 33%, *p* = 0.00079), and tumors of Grade Groups 1 and 2 (62% vs. 26%, *p* = 0.008794). SPINK1 overexpression was more in African American men (68% vs. 26%, *p* = 0.00008), in tumors with high tumor volume (> 20%) and with anterior located tumors. *ETV1* and *ETV4* demonstrated rare overexpression in these tumors, particularly in the higher-grade tumors. This study expands the knowledge of the clonal evolution of multifocal cancer in young patients and support differences in relation to racial background and genetics of prostate cancer.

## INTRODUCTION

Early-onset prostate cancer, defined as prostate cancer diagnosed at age ≤ 55 years, has shown different etiologic and clinical perspective from prostate cancer in older men (1). Ethnic, familial, and genetic factors play a significant role in early onset prostate cancer (2-5). However, the biological behavior of prostate cancers detected at a young age has not been adequately studied, and the genetic alterations are largely unknown. With the current molecular subtyping of prostate cancer, several important genetic alterations have been identified in prostate cancer based on cohorts from older men. However, the expression profiles of these genetic alterations in early-onset prostate cancer has not been well studied.

ETS (erythroblast transformation-specific) family gene fusions are commonly seen in prostate cancer [6]. *ERG* (ETS-related gene) is more frequently overexpressed in prostate cancer, followed by *ETV1* (ETS Variant 1), *ETV4* (ETS Variant 4), and *ETV5* (ETS Variant 5) (6). *SPINK1* (serine protease inhibitor Kazal-type 1) overexpression has also been described in prostate cancer, mainly in ERG negative prostate cancer, but its prognostic value is controversial (7). All these alterations are generally mutually exclusive in prostate cancer, but their expression profiles and probable roles in early-onset prostate cancer have not been thoroughly explored. Furthermore, conventional approaches use only the index/dominant nodule for evaluation. Given that approximately 60%-90% of prostate cancer has the tendency to develop as multifocal tumors (7-15), unlike the conventional approaches, we for the first time, characterized the expression profiles of *ERG, SPINK1, ETV1* and *ETV4* in early onset prostate cancer in 2 ethnic groups by using dual immunohistochemical and dual RNA *in-situ* hybridization analysis using whole mount radical prostatectomy tissue slides. We identified the clonal molecular heterogeneity by the mutually exclusive pattern of expression of more than 1 molecular marker in distinct tumor foci, and attempted to understand its potential role in early onset prostate cancer, and shed light on diagnostic and prognostic perspectives, and potential therapeutic stratification.

## MATERIALS AND METHODS

### Study Design and Patient Selection

A total of 151 radical prostatectomy specimens from patients younger than 55 years from January 2001 to December 2016 were retrieved from the archives of the Department of Pathology and Laboratory Medicine in the Henry Ford Health System. Clinical data recorded included age, race, family history, preoperative prostate-specific antigen, and biochemical recurrence. All cases were independently reviewed by 2 of the study authors (ZL and NSG). Pathologic parameters noted from review of radical prostatectomy whole mount tissue included tumor location, tumor volume, dominant nodule, secondary nodule, Grade group, pathologic T stage, margin status, and lymph node status. One representative whole mount tissue section was selected for evaluation of selected markers by immunohistochemistry and RNA *in-situ* hybridization.

### Histological Evaluation

Tumor location (posterior aspect vs. anterior aspect) and tumor volume (≤ 5%; 6%-10%; 11%-20%; and > 20%) was assessed from whole mount slides. We defined dominant nodule based on the pathologic significance, being the largest tumor size if Gleason score was similar between foci, highest Gleason score, or shortest distance from resection margins, or positive resection margin. The less pathologically significant tumor nodules were assigned as secondary nodules, which included smaller tumor size, lower Gleason score, or further from resection margins. Grade group, pathologic T stage, resection margin status, and lymph node status were also recorded.

### Dual immunohistochemistry and dual RNA *in-situ* hybridization on whole mount radical prostatectomy slides

Formalin-fixed paraffin-embedded whole mount radical prostatectomy tissues were thin-sectioned (4 µm) and mounted on 2 × 3-inch macro slides (50 × 76 mm) and stained for immunohistochemistry using antibodies against ERG/SPINK1 and RNA *in-situ* hybridization for *ETV1* and *ETV4* as described. Expression of ERG, SPINK1, *ETV1*, and *ETV4* were recorded based on staining intensity, positive tumor location and relationship to the dominant nodule and secondary nodules.

**Dual RNA *in-situ* hybridization** was performed as described previously (16, 17) on 2 × 3-inch slides. We have standardized procedures for 2 × 3-inch slides. Slides were incubated at 60°C for 1 hour. Tissues were then deparaffinized by immersing in xylene twice for 5 minutes each with periodic agitation. The slides were then immersed in 100% ethanol twice for 3 minutes each with periodic agitation, then air-dried for 5 minutes. Tissues were circled using a PAP pen (Vector, H-4000), allowed to dry, and treated with H_2_O_2_ for 10 minutes. Slides were rinsed twice in distilled water, and then boiled in 1X Target Retrieval Solution for 15 minutes. Slides were rinsed twice in distilled water, and then treated with Protease Plus solution for 15 minutes at 40°C in a HybEZ Oven (Advanced Cell Diagnostics, Newark, CA 321710). H_2_O_2_, 1X Target Retrieval, and Protease Plus are included in the RNAscope Pretreatment kit (Advanced Cell Diagnostics, 310020). Slides were rinsed twice in distilled water, and then treated with both *ETV1* (Advanced Cell Diagnostics, 311411), and *ETV4* (Advanced Cell Diagnostics, 478571-C2) probes at a 50:1 ratio for 2 hours at 40°C in the HybEZ Oven. Slides were then washed in 1X Wash Buffer (Advanced Cell Diagnostics, 310091) twice for 2 minutes each. Slides were then stored overnight in a 5X saline-sodium citrate solution. The next day, slides were again washed in 1X Wash Buffer twice for 2 minutes each. Slides were then treated with Amp 1 solution for 30 minutes, Amp 2 solution for 15 minutes, Amp 3 solution for 30 minutes, and Amp 4 solution for 15 minutes, all at 40°C in the HybEZ oven with 2 washes in 1X Wash Buffer for 2 minutes each after each step. Slides were then treated with Amp 5 solution for 30 minutes and Amp 6 solution for 15 minutes at room temperature in a humidity chamber with 2 washes in 1X Wash Buffer for 2 minutes each after each step. Red color was developed by adding a 1:60 solution of Fast Red B: Fast Red A to each slide and incubating for 10 minutes. Slides were washed in 1X Wash Buffer twice for 2 minutes each, then treated with Amp 7 solution for 15 minutes and Amp 8 solution for 30 minutes at 40° C in the HybEZ oven with 2 washes in 1X Wash Buffer for 2 minutes each after each step. Slides were then treated with Amp 9 solution for 30 minutes and Amp 10 solution for 15 minutes at room temperature in a humidity chamber with 2 washes in 1X Wash Buffer for 2 minutes each after each step. Brown color was developed by adding a solution of Betazoid DAB (1 drop DAB to 1 ml buffer; Biocare Medical, Pacheco, CA, BDB2004L) to each slide and incubating for 10 minutes. Amps 1-10 and Fast Red are included in the RNAscope 2.5 HD Duplex Detection Reagents (Advanced Cell Diagnostics, 322500). Slides were then rinsed 2 times in distilled water, then treated with EnVision FLEX Hematoxylin (Agilent DAKO, Santa Clara, CA, K800821-2) for 5 minutes. Slides were rinsed several times in distilled water, immersed in a 0.01% ammonium hydroxide solution, and then rinsed twice in distilled water. Slides were then dried completely. Slides were dipped in xylene approximately 15 times. EcoMount (Biocare Medical, EM897L) was added to each slide, which was then coverslipped.

**Dual ERG/SPINK1 Immunohistochemistry** was performed as follows. Slides were incubated at 60°C for at least 2 hours. Slides were then placed in a PT Link Pre-Treatment Module (Agilent DAKO, PT200) in high pH antigen retrieval solution (Agilent DAKO, K800421-2) and incubated at 90°C for 20 minutes. Slides were then washed in 1X EnVision FLEX Wash Buffer (Agilent DAKO, K800721-2) for 5 minutes. Slides were then treated with Peroxidazed 1 (Biocare Medical, PX968M) for 5 minutes and Background Punisher (Biocare Medical, BP974L) for 10 minutes with a wash of 1X EnVision FLEX Wash Buffer for 5 minutes after each step. Anti-ERG (EPR3864) rabbit monoclonal primary antibody diluted 1:50 (Abcam, Cambridge, MA, ab92513) and a mouse monoclonal against SPINK1 diluted 1:100 (Novus Biologicals, Centennial, CO, H00006690-M01) were added to each slide, which were then coverslipped with parafilm, placed in a humidifying chamber, and incubated overnight at 4°C. The next day, slides were washed in 1X EnVision Wash Buffer for 5 minutes and then incubated in Mach2 Doublestain 1 (Biocare Medical, MRCT523L) for 30 minutes at room temperature in a humidifying chamber. Slides were then rinsed in 1X EnVision Wash Buffer 3 times for 5 minutes each. Slides were then treated with a Ferangi Blue solution (1 drop to 2.5 ml buffer; Biocare Medical, FB813S) for 7 minutes, washed in 1X EnVision FLEX Wash Buffer for 5 minutes, and then treated with a Betazoid DAB solution (1 drop to 1 ml buffer) for 5 minutes. Slides were then rinsed 2 times in distilled water, then treated with EnVision FLEX Hematoxylin for 5 minutes. Slides were rinsed several times in distilled water, immersed in a 0.01% ammonium hydroxide solution, and then rinsed twice in distilled water. Slides were then dried completely. Slides were dipped in xylene approximately 15 times. EcoMount (Biocare Medical, EM897L) was added to each slide, which was then coverslipped.

### Statistical Analysis

The expression profiles were analyzed by using a chi-square statistics test.

## RESULTS

### Clinical Features

The clinical and histopathological features of the cohort used in the study are summarized in Table 1. A total of 151 patients younger than 55 years old with age ranging from 34 to 55 years (mean 49) were included in this study. Of the 151, 68 (45%) and 61 (40%) were Caucasian and African Americans, respectively. A total of 65 (43%) patients, of whom 29 were Caucasians and 29 were African Americans, had a positive family history of prostate cancer. Preoperative prostate-specific antigen ranged from 0.3 to 52.7 ng/ml (mean 7.04). Post-prostatectomy follow-up period ranged from 1 to 123.7 months (mean 30.3 months). Biochemical recurrence was observed in 8 (5%) patients, which included 3 African Americans and 2 Caucasians.

**Table 1:** The clinical and h istopathological features of the study cohort.

| Clinicopathologic variable | Whole cohort (n=151) | Caucasian (n=68) | African (n=61) |
| --- | --- | --- | --- |
| Age | 34-55 (mean 49) | 39-55 (mean 50) | 34-55 (mean 50) |
| Family history |  |  |  |
| positive | 65 | 29 | 29 |
| negative | 75 | 35 | 31 |
| unknown | 11 | 4 | 1 |
| Pre-operative PSA | 0.3-52.7 (mean 7.04) | 0.3-51.2 (mean 7.36) | 1.4-52.7 (mean 7.25) |
| Follow up months | 1-123.7 (mean 30.3) | 1-123.7 (mean 22.2) | 1-112.9 (mean 41.3) |
| Biochemical recurrence |  |  |  |
| positive | 8 | 2 | 3 |
| negative | 142 | 66 | 58 |
| unknown | 1 |  |  |
| Number of tumor nodules |  |  |  |
| 1 | 68 | 36* | 17* |
| 2 | 34 | 17 | 14 |
| 3 | 30 | 11 | 16 |
| 4 | 11 | 2 | 8 |
| 5 | 7 | 1 | 6 |
| 6 | 1 | 1 |  |
| Tumor location: |  |  |  |
| posterior or posterolateral peripheral zone (P) | 82 | 38 | 29 |
| anterior or anterolateral prostate (A) | 14 | 4 | 7 |
| A and P | 55 | 26 | 25 |
| Tumor volume |  |  |  |
| <6% | 48 | 24 | 16 |
| 6-10% | 41 | 24 | 13 |
| 11-20% | 38 | 14 | 18 |
| >20% | 24 | 6<sup>†</sup> | 14<sup>†</sup> |
| Gleason Score |  |  |  |
| GG-1 | 42 | 21 | 14 |
| GG-2 | 78 | 32 | 35 |
| GG-3 | 22 | 11 | 8 |
| GG-4 | 4 | 2 | 1 |
| GG-5 | 5 | 2 | 3 |
| Pathology Stage: pT2a |  |  |  |
| pT2a | 12 | 10<sup>‡</sup> | 0<sup>‡</sup> |
| pT2b | 3 | 1 | 2 |
| pT2c | 97 | 43 | 39 |
| pT3a | 29 | 12 | 13 |
| pT3b | 10 | 2 | 7 |
| Lymph node status |  |  |  |
| pN0 | 114 | 49 | 47 |
| pN1 | 7 | 2 | 4 |
| pNX | 30 | 17 | 10 |
| Tumor margins: positive |  |  |  |
| positive | 36 | 15 | 13 |
| negative | 115 | 53 | 48 |

\*p=0.0039, †p=0.0269, ‡p=0.0018,

### Histopathologic Features

A total of 313 prostate tumor foci were identified in 151 patients, ranging from 1 to 6 tumors per patient (mean 2). Over half of the 151 cases (n = 84, 56%), of which 44 were African American and 32 were Caucasian, had multifocal tumors. Statistical analysis showed that multifocal tumor was more frequent among young African American patients compared to Caucasian American patients (*p* = 0.0039). More than half of the cases (n = 82, 54%) had tumors predominantly located at the posterior or posterolateral peripheral aspect, whereas 14 cases (9%) were predominantly located in the anterior or anterolateral aspect of the prostate, and 55 cases (36%) had both posterior and anterior involvement. Tumor volumes were assessed by 5 tiers: 48 patients (32%) with tumor volumes of < 6%; 41 patients (27%) with tumor volumes of 6%-10%; 38 patients (25%) with tumor volumes of 11%-20%; and 24 patients (16%) with tumor volumes > 20%. Overall, 127 (84%) patients had low tumor volumes (≤ 20%). Comparison between Caucasian and African Americans showed that tumors with volumes greater than 20% were more common among African Americans (*p* = 0.0269). Grade groups were 42 (28%) patients with Grade Group 1 (3 + 3 = 6); 78 (52%) patients with Grade Group 2 (3 + 4 = 7); 22 (15%) patients with Grade Group 3 (4 + 3 = 7); 4 (3%) patients with Grade Group 4 (4 + 4 = 8); and 5 (3%) patients with Grade Group 5. Overall, 120 patients (79%) had low-grade tumors (Grade Groups 1 and 2). For pathologic T stage, tumors in 112 (74%) patients were organ confined (pT2); 29 (19%) patients had extraprostatic extension (pT3a); 10 (7%) patients had seminal vesicle involvement (pT3b). Statistical analysis revealed that pT2a tumor stage was more common among Caucasian Americans compared to African Americans (*p* = 0.0018). Lymph node status included 7 (5%) patients with positive lymph nodes (pN1), 114 (75%) with negative lymph nodes (pN0), and 30 (20%) with no lymph nodes submitted for evaluation (pNX). Only 36 patients (24%) showed positive tumor margins.

#### Evaluation of ERG/SPINK1/ETV1/ETV4 on whole mount radical prostatectomy tissue

Dual ERG/SPINK1 immunohistochemical staining and dual *ETV1*/*ETV4* RNA *in-situ* hybridization were performed on the whole mount radical prostatectomy on consecutive tissue slides. ERG overexpression was present in 85 (56%) cases, which included 48 (56%) Caucasians and 25 (29%) African Americans. SPINK1 overexpression was present in 61 (40%) cases, where 18 (30%) were Caucasian and 37 (61%) were African American. A total of 25 (17%) of the cases displayed co-expression of ERG and SPINK1 on independent tumor foci. Only a minority of patients had *ETV1* or *ETV4* overexpression, including 9 (6%) patients with *ETV1* overexpression where 2 (22%) were African American and 4 (44%) were Caucasian. Only 34 (2%) patients showed *ETV4* overexpression including 2 (67%) African Americans and 1 (33%) Caucasians.

ERG overexpression with strong diffuse nuclear staining pattern was observed in 85 (56%) of the cases. Frequent foci of high-grade prostatic intraepithelial neoplasia were identified with the same staining pattern adjacent to prostate cancer (Figure 1). The summary of clinicopathologic features of tumors with ERG overexpression are listed in Table 2. ERG was mainly overexpressed in the dominant tumor nodules 75/85 (88%), of which 60 showed staining exclusively in the dominant nodules, 15 cases in both dominant nodules and secondary nodules, and 10 cases in secondary nodules. Of the 85 cases, 76/85 (89%) were located at the posterior or posterolateral peripheral zone; 6/85 (7%) of cases were in the anterior/anterolateral location, and 3/85 (4%) displayed both posterior and anterior involvement. Most of the tumors 72/85 (85%) exhibited low tumor volumes (≤ 20%). This included 22 cases with tumor volumes of < 6%; 30 cases with tumor volumes of 6%-10%; and 20 cases with tumor volumes of 11%-20%. By contrast, 13 patients (15%) had tumor volumes > 20%. Most cases 74/85 (87%) were Grade Groups 1 and 2, of which 28/85 (33%) were Grade Group 1 and 46/85 (54%) were Grade Group 2. In contrast, 7/85 (8%) cases were Grade Group 3; 1/85 (1%) case was Grade Group 4; and 3/85 (4%) cases were Grade Group 5. Most cases, 72/85 (85%), were organ-confined (pT2), and a minority had extraprostatic extension, including 10 cases of pT3a and 3 cases of pT3b. Resection margins were positive in 16/85 (19%). Lymph nodes were positive in 2 cases (pN1), negative in 64 cases (pN0), and 19 cases with no lymph nodes resection performed (pNx). Three cases 3/85 (4%) had biochemical recurrence. In comparison to ERG negative prostate cancer, tumors with ERG overexpression were more commonly identified in patients younger than 45 years, 32/42 (76%), than older patients (older than 45 years but younger than 55 years, 53/109 [49%]; *p* = 0.002213). Tumors with ERG overexpression were more frequently identified in Caucasian men, 48/68 (71%), than those of African American men, 25/61 (41%, *p* = 0.000707). ERG overexpression was more frequently identified in tumors of Grade Groups 1 and 2 (62% vs. 35%, *p* = 0.008794). In addition, ERG overexpression was more commonly identified in organ-confined tumors, 72/112 (64%), than those tumors with extraprostatic extension, 13/40 (32.5%; *p* = 0.00079) (Table 2).

**Table 2:** The summary of clinicopathologic features of tumors with ERG overexpression.

| Clinicopathologic variable | ERG |  | P-value | SPINK1 |  | P-value |
| --- | --- | --- | --- | --- | --- | --- |
|  | Positive | Negative |  | Positive | Negative |  |
| Pre-operative PSA | 0.3 to 21.1 (6.26) | 1.4 to 52.7 (8.1) |  | 1.1 to 51.2 (6.84) | 0.3 to 52.7 (7.18) |  |
| Age: ≤45: 42(28%) | 32 | 10 |  | 14 | 28 |  |
| 46-55: 109 (72%) | 53 | 56 | 0.002213 | 47 | 62 | 0.27216 |
| Race: African American : 61 (40%) | 25 | 36 |  | 37 | 24 |  |
| Caucasian American : 68 (45%) | 48 | 20 | 0.000707 | 18 | 50 | 0.000089 |
| Family history: positive | 36 | 29 |  | 29 | 36 |  |
| negative | 41 | 34 | 0.932135 | 29 | 46 | 0.476085 |
| Pathology Stage: ≤ pT2:113 (74%) | 72 | 40 |  | 43 | 69 |  |
| ≥pT3a:40 (27%) | 13 | 26 | 0.00079 | 18 | 21 | 0.394953 |
| Tumor volume: <6% | 22 | 20 |  | 12 | 30 |  |
| 6-10% | 30 | 12 |  | 14 | 28 |  |
| 11-20% | 20 | 21 |  | 18 | 23 |  |
| >20% | 13 | 13 | 0.477193 | 17 | 9 | 0.004319 |
| Gleason Grade Group-1 | 28 | 14 |  | 17 | 25 |  |
| Gleason Grade Group-2 | 46 | 32 |  | 31 | 47 |  |
| Gleason Grade Group-3 | 7 | 15 |  | 11 | 11 |  |
| Gleason Grade Group-4 | 1 | 3 |  | 0 | 4 |  |
| Gleason Grade Group-5 | 3 | 2 | 0.008794 | 2 | 3 | 0.844784 |
| Dominant nodule | 60 | N/A |  | 32 | N/A |  |
| Secondary nodule | 10 |  |  | 17 |  |  |
| Dominant and secondary nodule | 15 |  |  | 12 |  | 0.038977 |
| Tumor Location : Posterior | 76 | N/A |  | 34 | N/A |  |
| Anterior | 6 |  |  | 22 |  |  |
| Posterior/anterior | 3 |  |  | 5 |  | .000011 |

**Figure 1:**
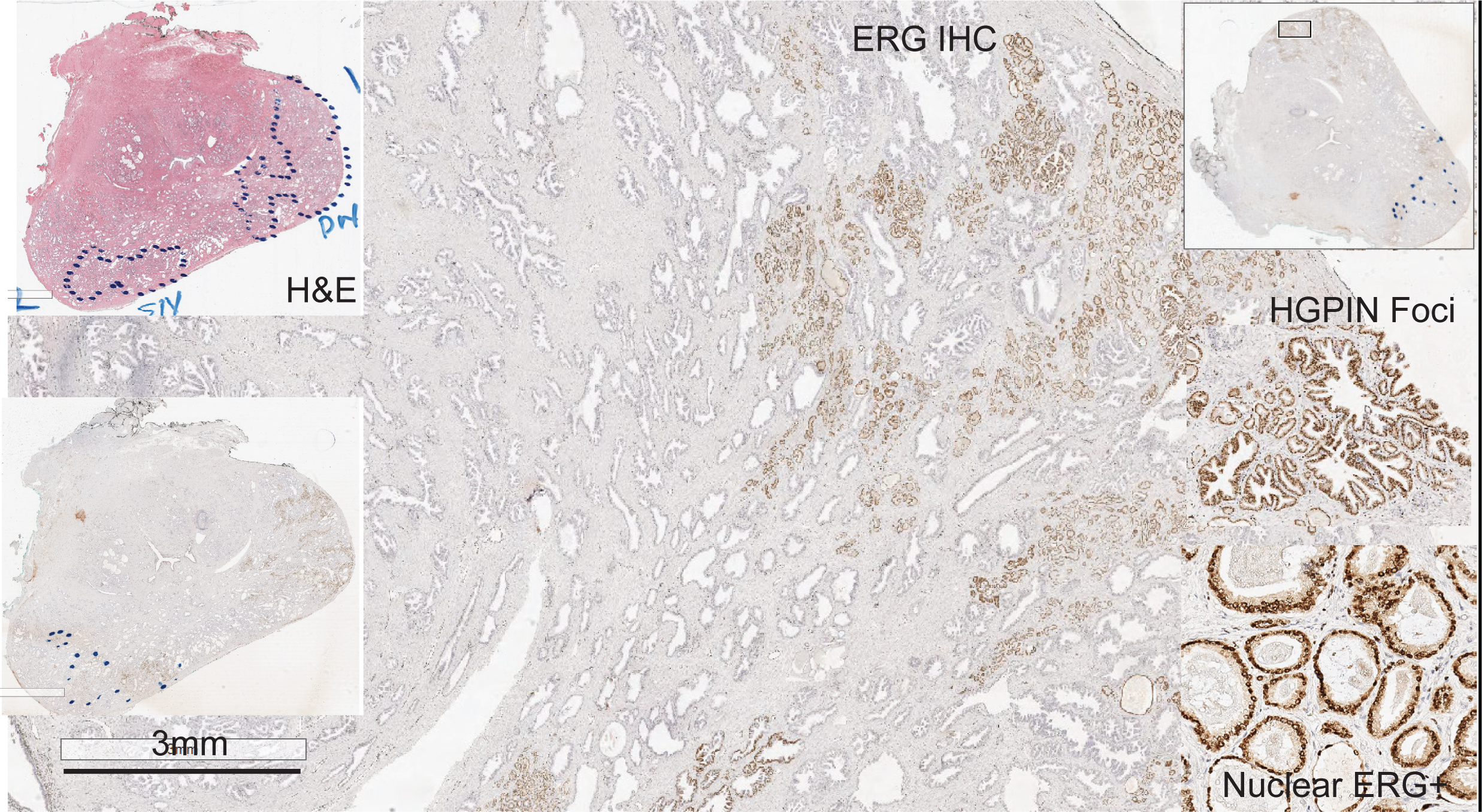
Dual IHC for ERG/SPINK1 on a whole mount prostatectomy tissue. ERG, strong nuclear positivity is seen in the dominant tumor and adjacent HGPIN and atrophic prostatic glands. Insets show the hematoxylin and eosin whole mount image marked with tumor foci (dotted circle, top left) and the high magnification view of ERG positive neoplastic glands with adjacent HGPIN. Abbreviations: HGPIN, high-grade prostatic intraepithelial neoplasia.

In our cohort, 61/151 (40%) cases exhibited SPINK1 overexpression with cytoplasmic and membranous staining pattern (Figure 2). SPINK1 heterogeneity, such as heterogeneous staining intensity between different tumor foci or within the same tumor focus, was noted. In addition, frequent foci of high-grade prostatic intraepithelial neoplasia were also highlighted by SPINK1 with the same staining pattern. The summary of clinicopathologic features of tumor with SPINK1 overexpression are listed in Table 2.

**Figure 2:**
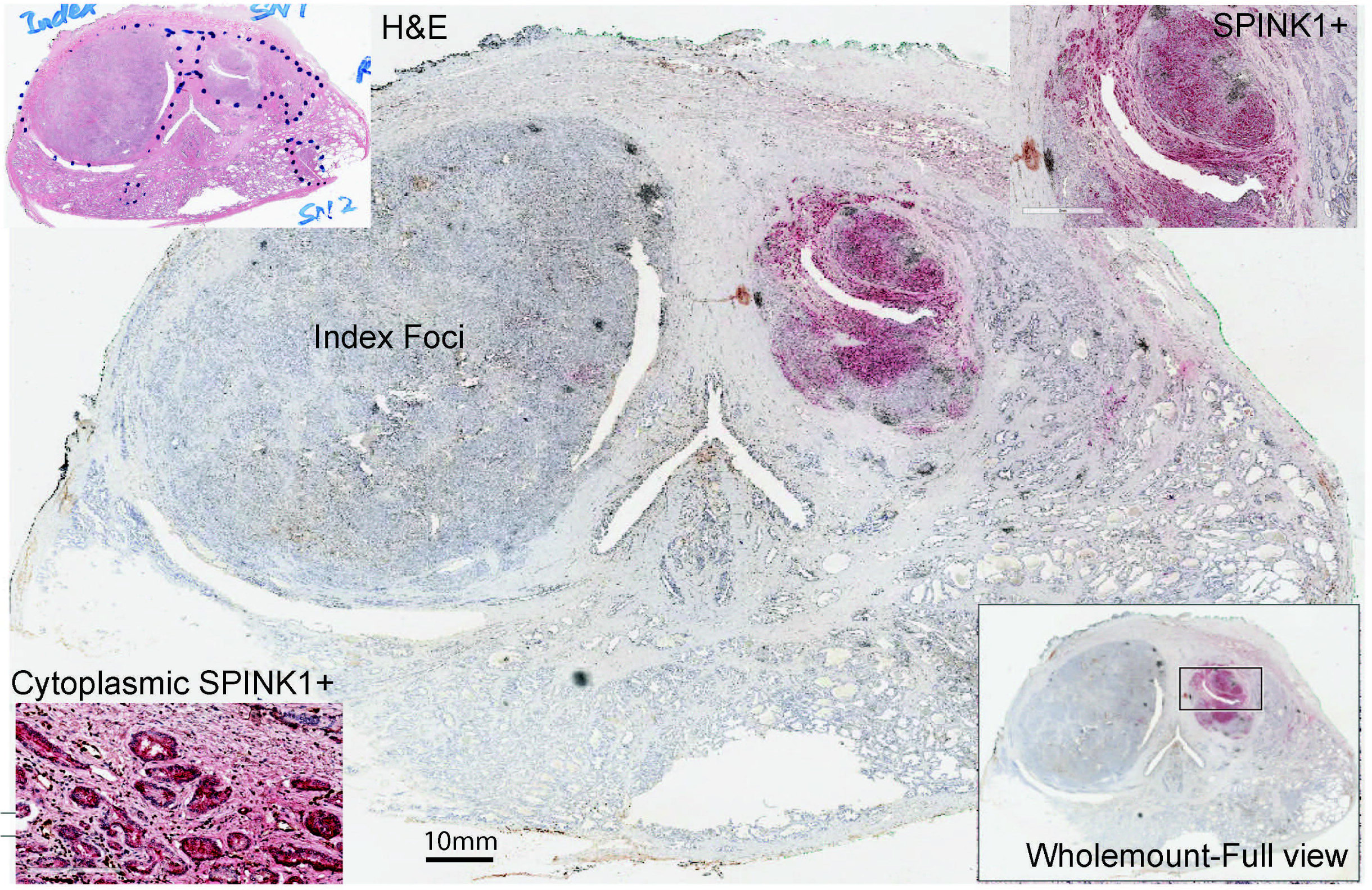
Dual IHC for ERG/SPINK1 on a whole mount prostatectomy tissue. **SPINK1,** strong cytoplasmic and membranous positivity is seen in a secondary tumor nodule while the index/dominant tumor nodule is negative for both ERG and SPINK1expression. Insets show the hematoxylin and eosin whole mount image (top left), high magnification view for SPINK1 positive tumor nodule (top right) and strong cytoplasmic staining of SPINK1 (bottom left) in tumor foci. ERG positivity is observed as brown nuclear staining in endothelial cells in blood vessels (internal control).

Of the 61 patients, 44 (72%) showed SPINK1 overexpression in dominant nodules, 17/61 (28%) were positive in secondary nodules, and 12/61 (20%) showed both dominant and secondary nodules expression. Over half of tumors, 34/61 (56%), involved the posterior or posterolateral peripheral zone; one-third, 22/61 (36%), of tumors involved the anterior or anterolateral, and 5/61 tumors showed both posterior and anterior involvement. The summary of clinicopathologic features of SPINK1 overexpression in anterior/anterolateral predominant tumors are listed in Table 3. About two-thirds of these patients, 44/61 (72%), had tumor volumes of < 20%, including 12 patients with tumor volumes of < 6%; 14 patients with tumor volumes of 6%-10%; 18 patients with tumor volumes of 11%-20%. In contrast, 17 tumors had volumes > 20%. Four fifth of tumors, 48/61 (79%), were low Grade Groups 1 and 2, including 17/61 with Gleason score 3 + 3 = 6 (Grade Group 1), 31/61 with Gleason score 3 + 4 = 7 (Grade Group 2); 11/61 with Gleason score 4 + 3 = 7 (Grade Group 3); and 2/61 were Grade Group 5. Majority of tumors 43/61 (70%) were organ-confined (pT2), and a minority of tumors showed extraprostatic extension, including 12 pT3a and 6 pT3b and 17/61 (28%) tumors with positive resection margins. Lymph node status included 2 patients with pN1, 47 with pN0, and 12 with pNX. Three patients, 3/61 (5%), experienced biochemical recurrence.

**Table 3:**
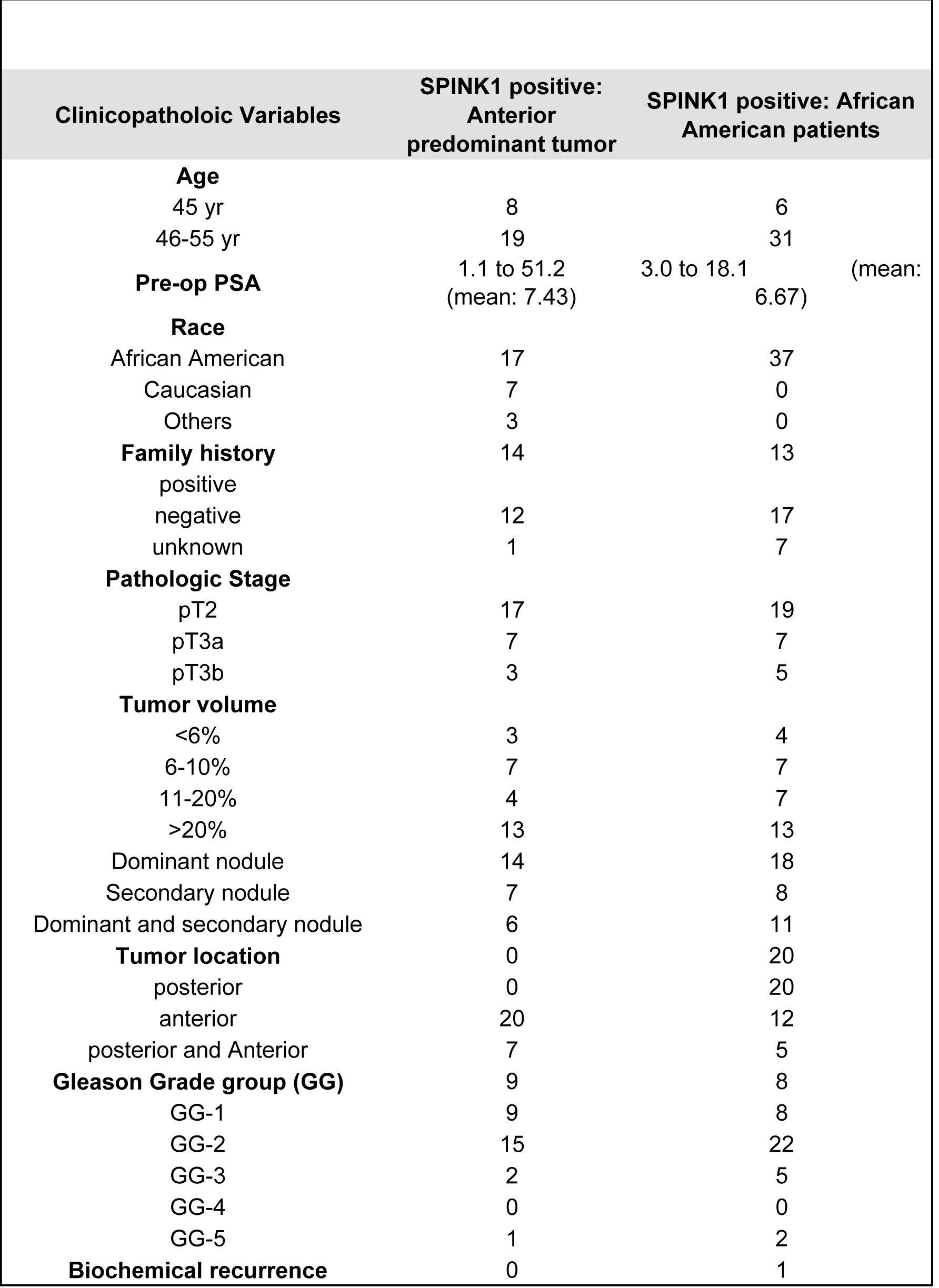
The summary of clinicopathologic features of SPINK1 overexpression in anterior/anterolateral predominant tumors

In comparison to SPINK1 negative tumors, SPINK1 overexpression was more often seen in the tumors of African American men, 37/61 (61%), than those from Caucasian men, 18/68 (26%; *p* = 0.000089). The summary of clinicopathologic features of SPINK1 positive prostate cancer in African American patients is listed in Table 3. SPINK1 overexpression was more frequently identified in tumors with high tumor volume (tumor volume > 20%) than those with tumor volume < 20% (17/26 [65%] vs. 44/125 [35%]; *p* = 0.004319). There was no association with other pathologic parameters (Table 2).

In 25/151 (17%) patients with multifocal tumors concomitant overexpression of ERG and SPINK1 (Figure 3) was observed. Of the 25 tumors, 7 displayed concomitant positivity for ERG and SPINK1 in the same nodule but not in the same tumor cells (including 7 dominant nodules and 2 secondary nodules), and 18 in different nodules of prostate cancer indicating intra- and inter-tumor molecular heterogeneity. Most of the tumors, 19/25 (76%), were of tumor volumes ≤ 20%, including 6 with tumor volumes of < 6%, 6 with tumor volumes of 6%-10%, and 7 with tumor volumes of 11%-20% in contrast to 6 cases with tumor volumes > 20 %. Most of the cases (84%, 21/25) were Grade Groups 1 and 2, including 8/25 Gleason score 3 + 3 = 6 (Grade Group 1), 13/25 Gleason score 3 + 4 = 7 (Grade Group 2); and 3/25 Gleason score 4 + 3 = 7 (Grade Group 3). While 1/25 was Grade Group 5. Most tumors (88%, 22/25) were organ-confined (pT2), whereas 3 had extraprostatic extension, including 1 pT3a and 2 pT3b. Positive resection margins were present in 6/25 tumors. Lymph nodes were positive in 1 patient (pN1), whereas 20 were negative (pN0) and 4 were pNX.

**Figure 3:**
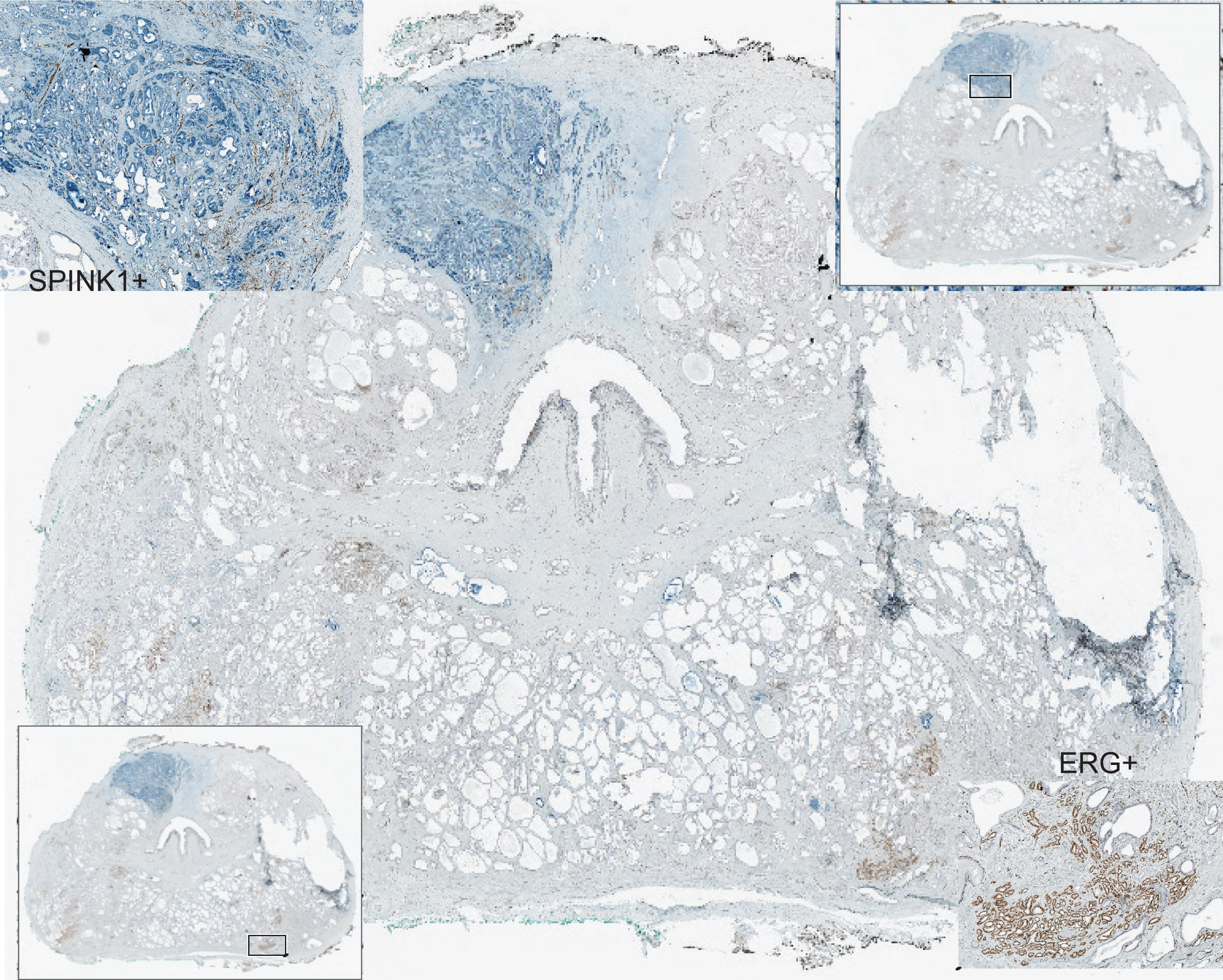
Dual IHC for ERG/SPINK1 on a whole mount prostatectomy tissue. Multiple independent tumor foci showing mutually exclusive expression of ERG (stained in brown) and SPINK1 (stained in blue). Insets showing the high magnification view for ERG (bottom right) and SPINK1 (top left) expression. There are multiple secondary tumor foci separated by benign glands/stroma showing ERG which indicates inter tumor heterogeneity with independent clonal origin of the tumors.

In our study cohort, 9/149 (6%) patients showed strong *ETV1* expression (2 African American and 4 Caucasians), which included 2 patients with tumor volumes > 20% and 7 with tumor volumes < 20% (Figure 4). Most cases were Grade Groups 2 (6/9) and 3 (2/9), 1/9 was Grade Group 1. There were 5/9 (55%) patients with pT3 stage and 4/5 of these showed positive margins. One patient had positive lymph nodes. Biochemical recurrence was not encountered in any of these cases. In the conjunction with ERG and SPINK1 status, 5 cases showed coexpression of ERG and SPINK1 in the different tumor foci, including 2 cases ERG positive/SPINK1 negative, 3 cases ERG negative/SPINK1 positive. Four patients were both ERG and SPINK1 negative.

**Figure 4:**
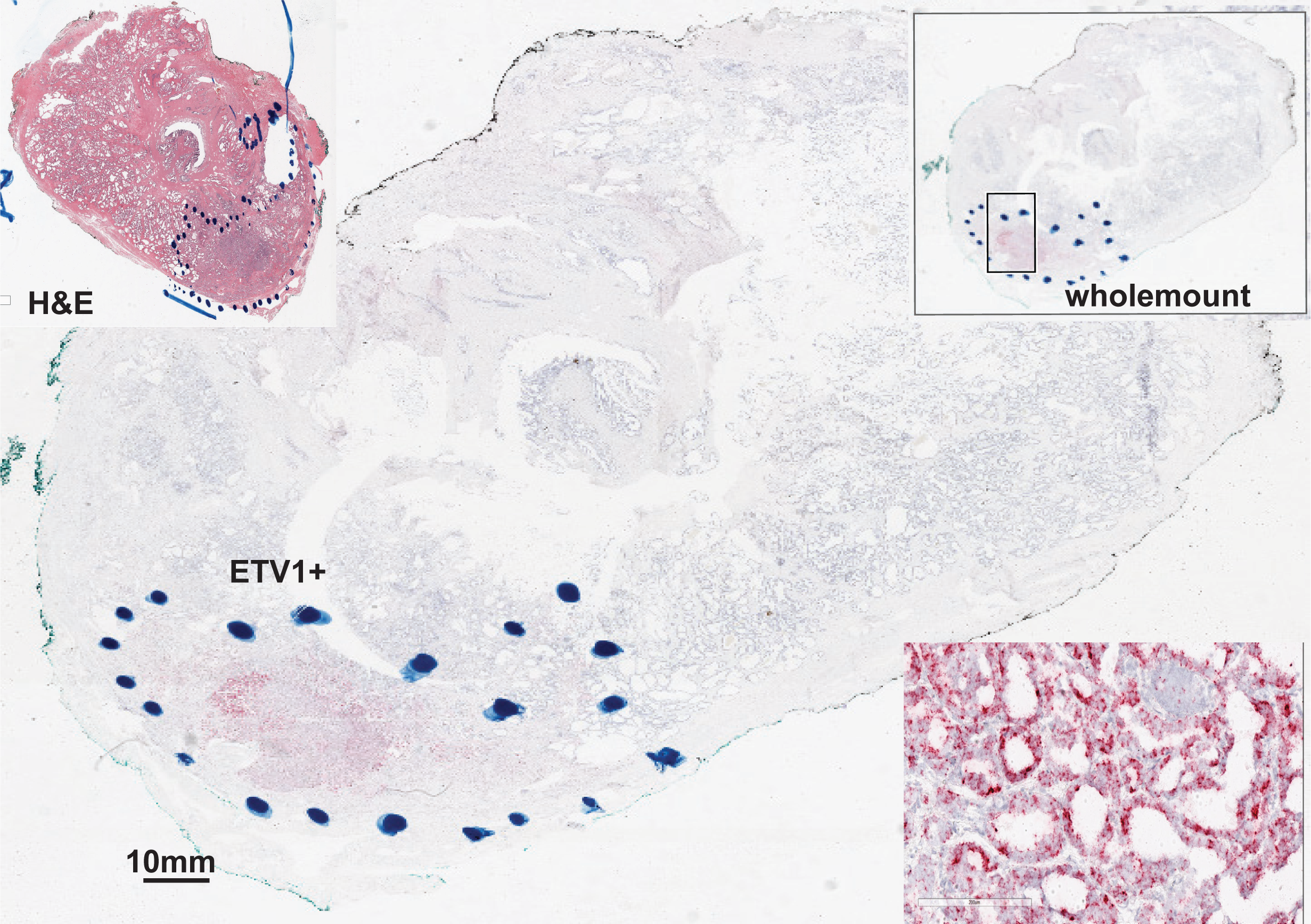
ETV1 RNA in situ hybridization on a whole mount prostatectomy tissue. The dominant tumor focus shows strong cytoplasmic and membranous expression of ETV1 whereas the secondary tumor focus is negative for ETV1 expression. Insets show the whole mount image of the H&E stain (top left) and ETV1 ISH (top right), the high magnification view of tumor focus with ETV1 expression (bottom right).

*ETV4* overexpression was detected in 3/141 (2%) of patients (two African American and one Caucasian American) of which two cases with tumor volumes of > 20% and one case with a tumor volume of 11%-20% (Figure 5). One tumor was Grade Group 5, one Grade Group 2, and 1 was Grade Group 1. Three demonstrated high pathologic state (pT3) with positive resection margins. One case showed positive lymph node involvement. None of these cases showed biochemical recurrence. As for the concurrent ERG and SPINK1 status, 1 was ERG+/SPINK1-, 1 was ERG-/SPINK1+, 1 was negative for both.

**Figure 5:**
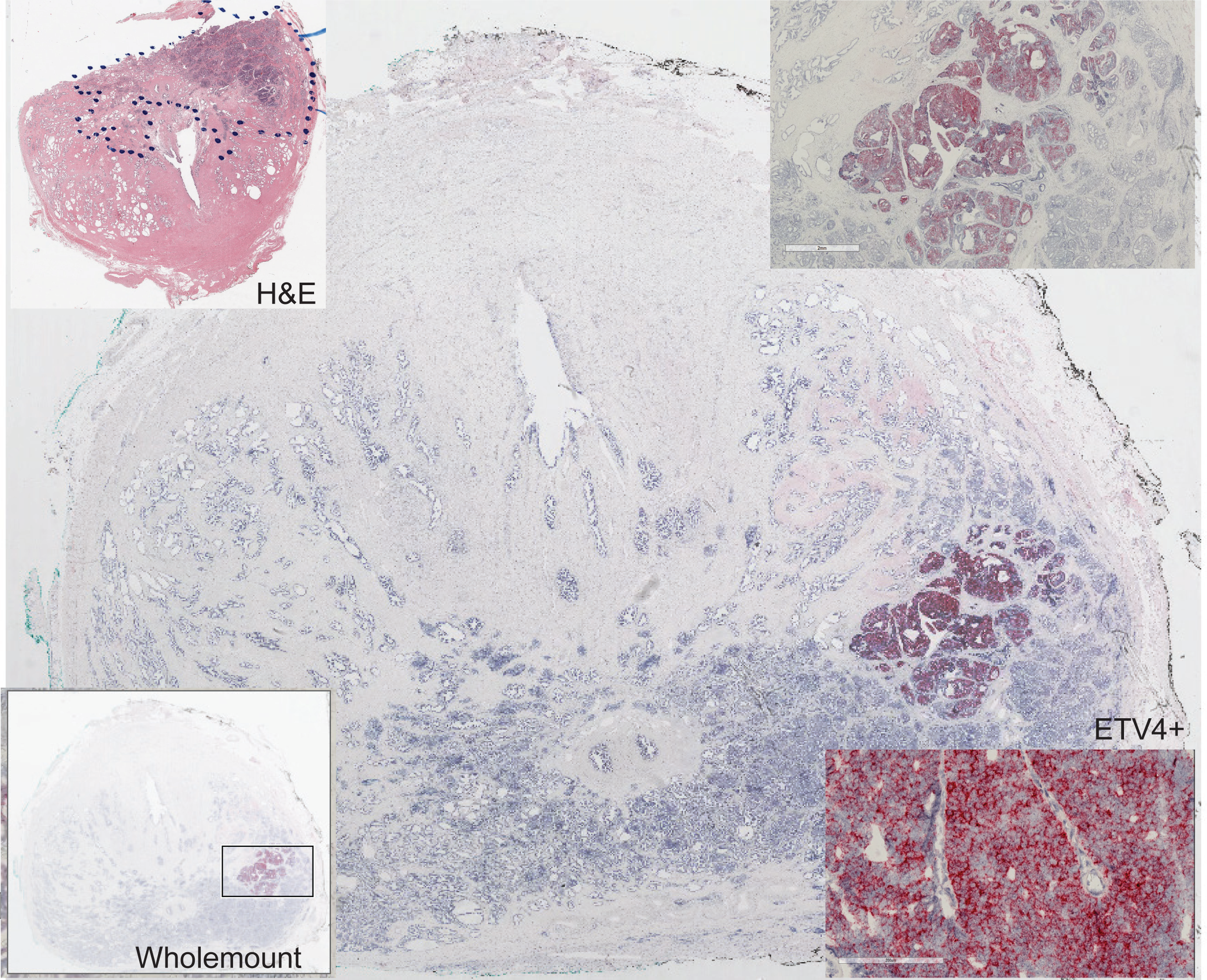
ETV4 RNA in situ hybridization on a whole mount tissue. The subset of dominant tumor nodule shows strong cytoplasmic and membranous expression of ETV4; in contrast, the adjacent subset of neoplastic glands of the dominant tumor nodule shows no expression of ETV4. This feature demonstrates the intra tumor heterogeneity. Insets show the H&E image of whole mount tissue (top left) and the high magnification view of tumor focus with ETV4 expression (bottom left)

Overall, in our cohort, 136 cases had an evaluable expression profile of all 4 biomarkers. Of these, 38 (28%, 38/136) exhibited more than 1 biomarker overexpression, 47 (34%) were only ERG positive, 32 (24%) were only SPINK1 positive, 3 (2%) were *ETV1* positive only, and 16 (12%) were negative for all 4 biomarkers (Figure 6). In further correlative analysis of expression of ERG and SPINK1 in dominant and secondary foci with Gleason grade group, a total of 313 prostate tumor foci were identified in 151 patients, which included 151 dominant nodules and 162 secondary nodules. Of dominant nodules, most (79%, 119/151) were Grade Groups 1 and 2, 48% (73/151) were ERG positive/SPINK1 negative, including 32% (23/73) Grade Group 1, 56% (41/73) Grade Group 2, 7% Grade Group 3, 2% Grade Group 4, and 4% Grade Group 5. ERG negative/SPINK1 positive was found in 25% (37/151), including 24% of cases of Grade Group 1, 54% of Grade Group 2, 19% of Grade Group 3, and 3% of Grade Group 5. Only a few cases (5%) were both ERG and SPINK1 positive, with Grade Groups 3 and lower. About one-quarter of cases (23%) were both ERG and SPINK1 negative, in which 24% were Grade Group 1, 38% were Grade Group 2, 26% were Grade Group 3, 9% were Grade Group 4, and 3% were Grade Group 5. Of secondary nodules, most (89%, 144/162) were Grade Group 1, including 48% (77/162) being both ERG and SPINK1 negative. Of secondary nodules, 22% were ERG positive/SPINK1 negative, 29% were ERG negative/SPINK1 positive, and 1% was both ERG and SPINK1 positive (Table 4).

**Table 4:** ERG/SPINK1 expression in dominat and secondary tumor nodules.

| **DN (151)** | **GG-1 (42)** | **GG-2 (77)** | **GG1-2** | **GG-3 (23)** | **GG 4 (4)** | **GG5(5)** | **GG 3-5** | **P-value** |
| --- | --- | --- | --- | --- | --- | --- | --- | --- |
| ERG+/SPINK1- (73) | 23 (32%) | 41 (56%) | 54 | 5 (7%) | 1(1%) | 3 (4%) | 9 |  |
| ERG-/SPINK1+ (37) | 9 (24%) | 20 (54%) | 29 | 7 (19%) | 0 | 1 (3%) | 8 |  |
| ERG+/SPINK1+ (7) | 2 (29%) | 3 (43%) | 5 | 2 (29%) | 0 | 0 | 2 | .676709 |
| ERG-/SPINK1- (34) | 8 (24%) | 13 (38%) | 21 | 9 (26%) | 3 (9%) | 1 (3%) | 4 | >0.05 |
| **SNs (162)** | **GG-1 (144)** | **GG-2(18)** | **GG1-2** | **GG-3 (0)** | **GG-4 (0)** | **GG-5 (0)** | **GG 3-5** | **P-value** |
| ERG+/SPINK1- (35) | 31(89%) | 4 (11%) | 35 | 0 | 0 | 0 | 0 |  |
| ERG-/SPINK1+ (48) | 38(79%) | 10(21%) | 48 | 0 | 0 | 0 | 0 |  |
| ERG+/SPINK1+ (2) | 2(100%) | 0 | 2 | 0 | 0 | 0 | 0 |  |
| ERG-/SPINK1- (77) | 73(95%) | 4(5%) | 77 | 0 | 0 | 0 | 0 |  |

**Figure 6:**
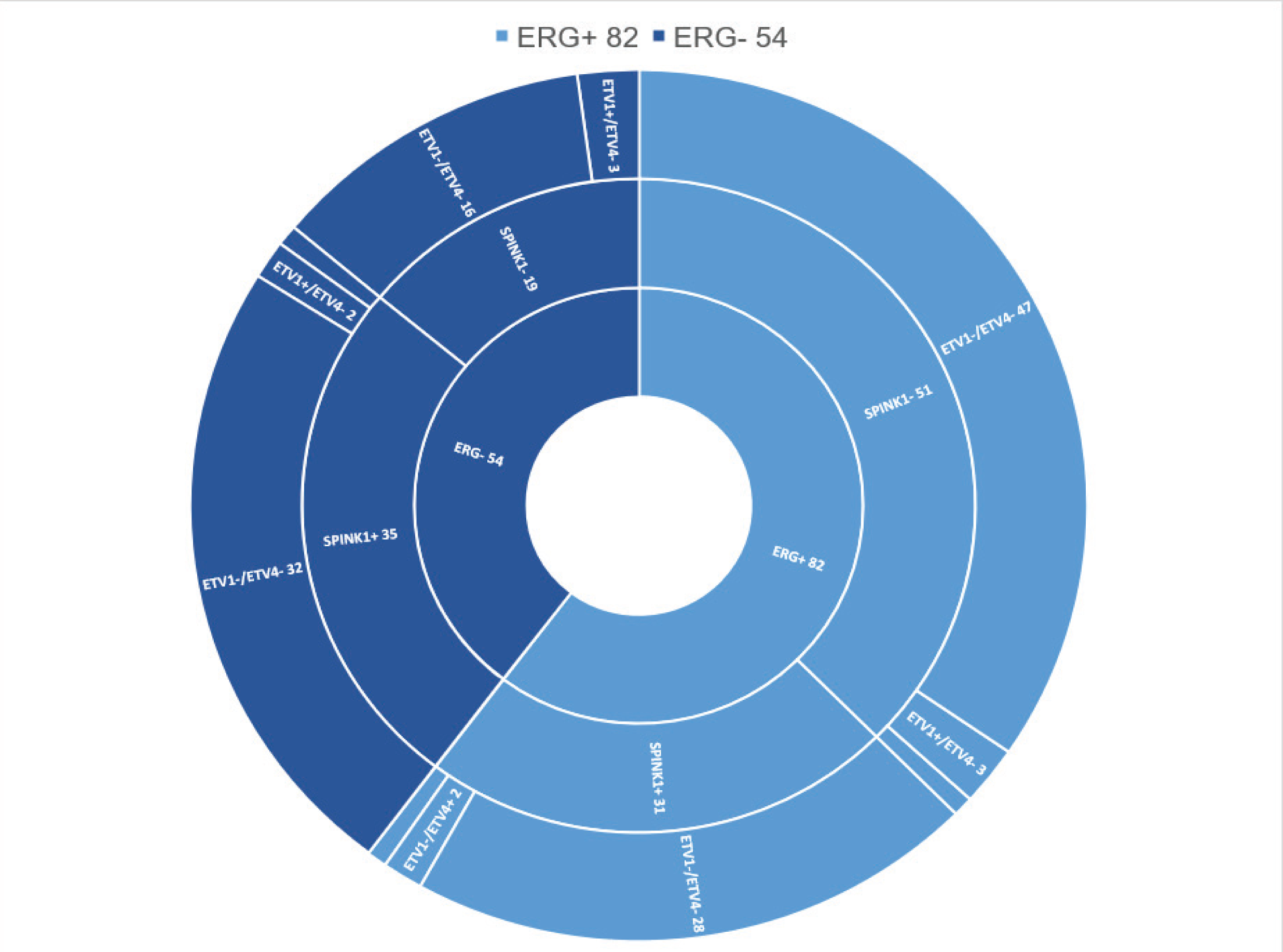
Sunburst chart showing the summary of distribution of four biomarkers (ERG, SPINK1, *ETV1*, and *ETV4*) mutually exclusive expression on whole mount radical prostatectomy tissue in the young men with prostate cancer, highlighting ERG positive and ERG negative tumors with SPINK1, *ETV1* and *ETV4* expression status.

In comparison between ERG positive dominant nodules and SPINK1 positive dominant nodules, ERG positive dominant nodules tend to be more common in patients younger than 45 years old (*p* = 0.013724); SPINK1 positive dominant nodules tend to be higher pathologic stage (≥ pT3) (*p* = 0.013308), higher tumor volume (> 20%; *p* = 0.007289), and anterior located tumors (*p* = 0.000313). There are no significant differences found in other parameters. In addition, there is no significant difference between Gleason grade group and ERG or SPINK1 overexpression status (*p* = 0.676709) (Table 5).

**Table 5:**
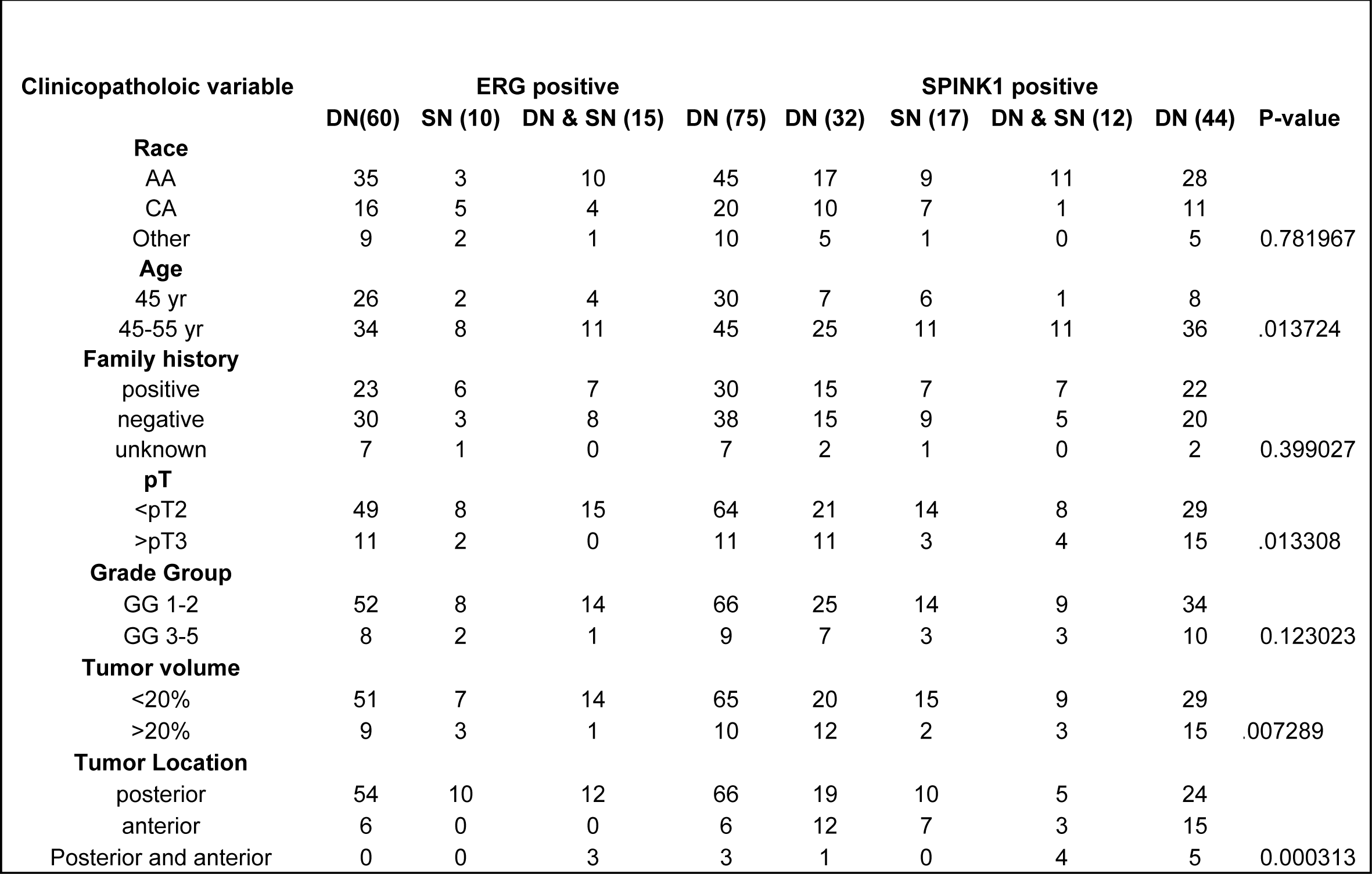
ERG/SPINK1 expression in dominat and secondary nodule and association with gleason group and other clinical parameters.

## DISCUSSION

For the first time, we performed an evaluation of ETS genes and SPINK1 overexpression on whole mount radical prostatectomy tissues to evaluate the clonal heterogeneity in young men with prostate cancer, in a racial disparity perspective. The key observations are the mutually exclusive expression of more than 1 marker in independent tumor foci in multifocal tumors. We did not observe concomitant expression of either of these 4 markers studied in any tumor foci suggesting the independent clonal evolution of tumors with different driver molecular aberrations. Although morphological tumor heterogeneity has been well studied, we have demonstrated the molecular heterogeneity based on the well-characterized prostate cancer specific tumor markers. Based on these observations we have identified new molecular subtypes of prostate cancer. The implication of these observations, based on the clinical correlations as discussed below, have direct applications to evaluate the tumor molecular heterogeneity in biopsy specimens in order to make informed decisions on active surveillance.

The *TMPRSS2-ERG* fusion, the most common genetic alteration identified in prostate cancer, has been found to be an early event in prostate cancer (18, 19). In our study series, ERG overexpression was identified in 56% (85/151) of patients. ERG overexpression was more commonly identified in younger patients (≤ 45 years old) (*p* = 0.002213), Caucasian patients (*p* = 0.000707), organ-confined tumors (*p* = 0.00079), and lower Grade Groups 1 and 2 tumors (*p* = 0.008794). Similarly, studies from Steurer’s group in Germany and Schaefer’s group in Austria both demonstrate that ERG positive prostate cancers are strongly linked to young patient age in low-grade prostate cancer (20, 21).

SPINK1, also known as Serine peptidase inhibitor, Kazal type 1, is commonly overexpressed in SPOP-mutant/CHD1 deleted ETS-negative prostate cancers. It has been identified as the second-most common biomarker for subclassification of prostate cancers, with around 10% of overexpression in prostate cancer. It is generally considered mutually exclusive from *ERG* rearrangements (22, 23). In our cohort, SPINK1 demonstrate significant higher overexpression (40%) in young age men with prostate cancer, particularly, in African-American men (68% vs. 26%, *p* = 0.00008). Furthermore, SPINK1 overexpression is more frequently identified in prostate cancer with higher tumor volume (> 20%) (28% vs. 10%; *p* = 0.004319) and dominant tumor nodule with an overall high pathologic stage (≥ pT3) (p=0.013308. Studies from Powell et al. and Johnson et al. also demonstrate that African-American men are at a higher risk of being diagnosed with advanced stage prostate cancer at a young age, with SPINK1 highlighting prostate cancer with more rapid progression (24, 25). Therefore, ERG and SPINK1 overexpression status in conjunction with dominant nodules status may help with stratifying prostate cancer.

Interestingly, in our cohort, anteriorly located prostate tumors tend to be more frequently positive for SPINK1 compared to ERG (39% [27/69] vs. 13% [9/69]). Of 61 SPINK1 positive prostate cancers, 17/37 prostate cancers from African American men showed anterior involvement. It is well documented that prostate cancer from African American men have higher frequency of anterior involvement (26). Our observation suggests that SPINK1 might be a more specific biomarker for anterior zone located prostate cancer in African American patients, with a potential indicator for higher pathologic stage tumor.

Prostate cancer is well known to be characterized with clonal heterogeneity. In our cohort, 25/151 (17%) cases were concomitantly positive for ERG and SPINK1. Of these, 7 cases showed a merging growth pattern in the same tumor focus, potentially demonstrating the clonally heterogeneous nature of prostate cancer. *ETV1* and *ETV4* are commonly associated with more aggressive prostate cancer (27, 28). In our study, rare *ETV1* and *ETV4* overexpression were identified in early-onset prostate cancer, mostly associated with higher grade features, which further supports the overall nature of early-onset prostate cancer.

In summary, our study provides some evidence that early-onset prostate cancer tends to be low-grade, low-stage, and the majority behave in an indolent fashion with low rates of biochemical recurrence rates. A higher frequency of ERG expression was seen in younger aged patients (≤ 45 years old), Caucasian patients, organ-confined tumors, and lower Grade group (1 and 2). SPINK1 immunoreactivity is more often seen in tumors with higher tumor volume (> 20%) in African American patients with frequently anterior tumor labelling. ERG and SPINK1 co-expression was observed in 17% of cases within different regions of the same tumor or different tumors within the same prostate gland, which demonstrates the heterogeneous nature of prostate cancer. *ETV1/ETV4* generally displays very low positivity in young prostate cancer patients (≤ 55 years old) with the potential role for detection of more aggressive prostate cancer. A panel of biomarkers (ERG, SPINK1, and *ETV1/4*) may be a useful tool to stratify prostate cancer, especially in early onset prostate cancer, which can be used for guiding individual therapeutic options.

To our knowledge, this is the first study to report 4 common biomarkers of prostate cancer reflecting genetic alterations on early onset prostate cancer by use of whole mount prostate cancer tissue. It provides a resource of continued investigation into the molecular and biological heterogeneity of early-onset prostate cancer.

## ACKNOWLEDGEMENTS

We thank Natalia Draga for her assistance in the preparation of slides from whole mount prostatectomy blocks. This work was supported in part by a US Department of Defense grant W81XWH-16-1-0544 to NP.

## Disclosure/Conflict of Interest

All authors disclose no conflict of interest.

